# An aphrodisiac produced by *Vibrio fischeri* stimulates mating in the closest living relatives of animals

**DOI:** 10.1101/139022

**Authors:** Arielle Woznica, Joseph P Gerdt, Ryan E. Hulett, Jon Clardy, Nicole King

## Abstract

We serendipitously discovered that the marine bacterium *Vibrio fischeri* induces sexual reproduction in one of the closest living relatives of animals, the choanoflagellate *Salpingoeca rosetta*. Although bacteria influence everything from nutrition and metabolism to cell biology and development in eukaryotes, bacterial regulation of eukaryotic mating was unexpected. Here we show that a single *V. fischeri* protein, the previously uncharacterized EroS, fully recapitulates the aphrodisiac activity of live *V. fischeri*. EroS is a chondroitin lyase; although its substrate, chondroitin sulfate, was previously thought to be an animal synapomorphy, we demonstrate that *S. rosetta* produces chondroitin sulfate and thus extend the ancestry of this important glycosaminoglycan to the premetazoan era. Finally, we show that *V. fischeri*, purified EroS, and other bacterial chondroitin lyases induce *S. rosetta* mating at environmentally-relevant concentrations suggesting that bacterially-produced aphrodisiacs likely regulate choanoflagellate mating in nature.

## Introduction

Bacterial–eukaryotic interactions are ubiquitous, and the influences of bacteria on eukaryotes vary from subtle to profound. Yet, because eukaryotes are often associated with complex and unseen communities of bacteria, the breadth of eukaryotic biological processes regulated by bacteria and the underlying molecular dialogue often remain obscure. Nonetheless, studies of experimentally tractable host-microbe pairs have revealed a growing number of examples in which bacteria regulate eukaryotic cell biology and development, in some cases using molecular cues that mediate pathogenesis in other contexts (Koropatnick et al., 2004).

One of the closest living relatives of animals, the marine choanoflagellate *S. rosetta*, has emerged as an attractive model for uncovering bacterial cues that regulate eukaryotic development. Like all choanoflagellates, *S. rosetta* survives by eating bacteria (Dayel and King, 2014; Leadbeater, 2015). However, interactions between *S. rosetta* and bacteria extend far beyond those of predator and prey. In prior work, we demonstrated that the developmental switch triggering the formation of multicellular “rosettes” from a single founding cell (Fairclough et al., 2010) is regulated by specific lipids produced by the environmental bacterium *Algoriphagus machipongonensis* (Alegado et al., 2012; Cantley et al., 2016; Woznica et al., 2016). Rosette development is one of at least six different developmental switches in the sexual and asexual phases of *S. rosetta*’s dynamic life history (Dayel et al., 2011; Levin and King, 2013), but until now was the only choanoflagellate process known to be regulated by bacterial cues.

We report here on our serendipitous discovery that sexual reproduction in *S. rosetta* is regulated by a secreted cue from the marine bacterium *Vibrio fischeri*.

## Results

### *S. rosetta* forms mating swarms upon exposure to *V. fischeri*

*V. fischeri* is perhaps best understood as a model for bacterial quorum sensing and as a symbiont required for the induction of light organ development in the squid, *Euprymna scolopes* (Mcfall-Ngai, 2014). Although *Vibrio* spp. are known symbionts, commensals, and pathogens of animals (Thompson et al., 2006), *V. fischeri* does not induce rosette development (Alegado et al., 2012) and was not previously known to influence any aspect of *S. rosetta* biology. We were therefore surprised to observe that the addition of live *V. fischeri* bacteria to a culture of single-celled, motile *S. rosetta* induced cells to gather rapidly into loose aggregates or “swarms,” each composed of between 2-50 cells (Figures 1A,B, Figure S1A, Supplemental Movie 1). This dynamic and previously unobserved swarming behavior began as early as 15 minutes after induction with *V. fischeri*, with individual *S. rosetta* cells often moving between swarms that periodically broke apart or merged with other swarms. In its timescale, mechanism, and outcome, swarming was clearly unrelated to the *Algoriphagus*-induced developmental process by which a single *S. rosetta* cell divides repeatedly to form a rosette (Fairclough et al., 2010).

**Figure 1.**
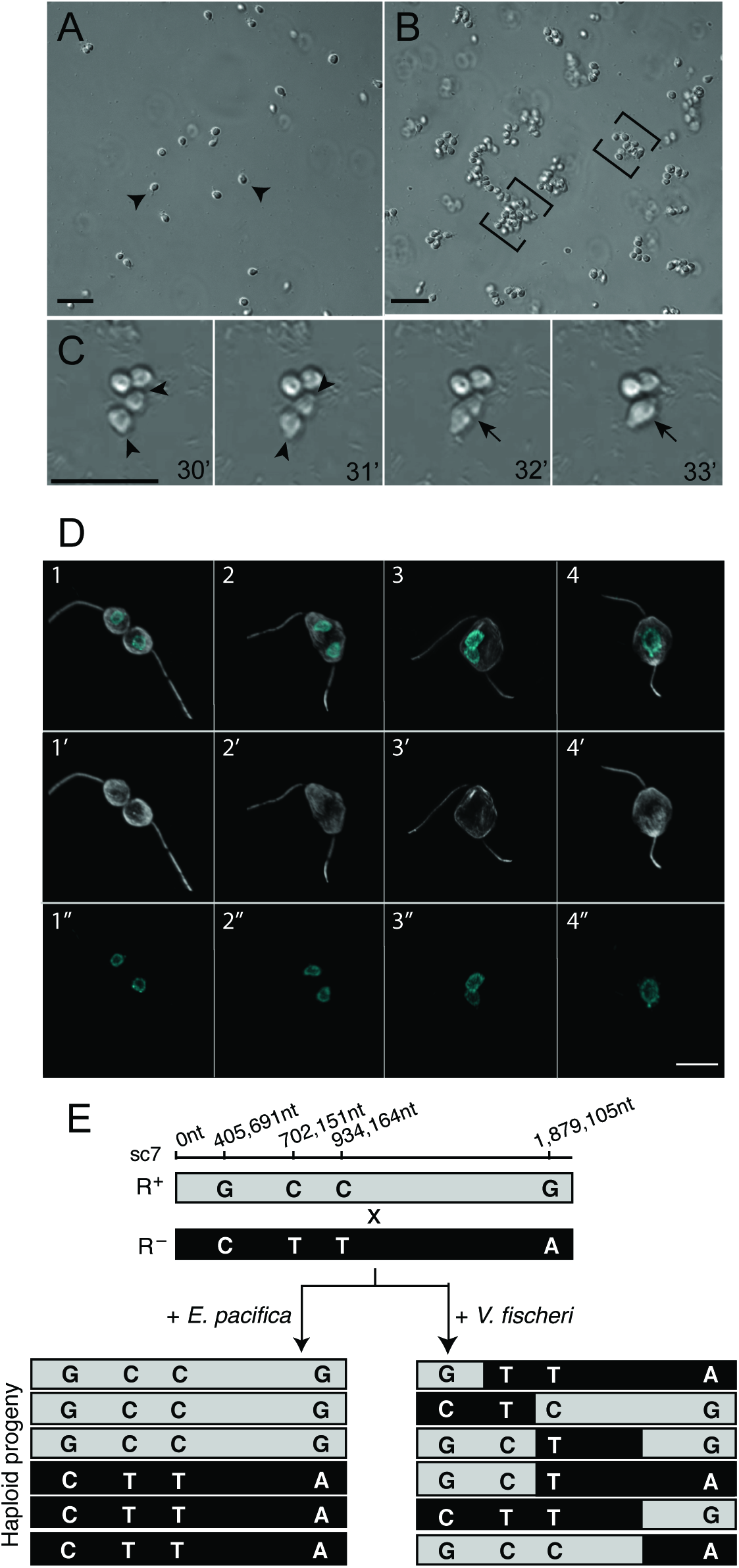
*V. fischeri* bacteria induce swarming and mating in the choanoflagellate, *S. rosetta*. **(A)** In the absence of *V. fischeri*, motile *S. rosetta* cells (arrowheads) are evenly dispersed. **(B)** Within 30 minutes of exposure to *V. fischeri, S. rosetta* motile cells aggregate into large swarms (brackets). Scale bar = 20µm. **(C)** *S. rosetta* cells within a swarm pair and fuse. Prior to fusion, cells reposition themselves such that their basal membranes are adjacent and their apical flagella point away (31’; arrowheads mark apical pole of unfused cells). Cell fusion takes only minutes, and occurs along the basal membrane (32’; indicated by arrow), resulting in a single, elongated cell (33’; indicated by arrow). Scale bar = 20µm. **(D)** Stages of cell and nuclear fusion in *S. rosetta* mating pairs. Haploid mating pairs are oriented with their basal poles (opposite the flagellum) touching (D1), and cell fusion proceeds along the basal membrane, resulting in a binucleated cell with two flagella (D2). Nuclei then congress towards the midline (D3), where the nuclei undergo nuclear fusion, resulting in a diploid cell (D4). Anti-tubulin antibody (D1’-4’; white) highlights the cell body and flagellum, and Hoechst (D1’’-4’’; cyan) highlights the nucleus. Scale bar = 5µm **(E)** Evidence for meiotic recombination in *S. rosetta* following exposure to *V. fischeri*. Two haploid, genotypically distinct *S. rosetta* strains [R+(grey shading) and R-(black shading)] were mixed in the presence of either *E. pacifica* conditioned media (EPCM) or *V. fischeri* conditioned media (VFCM) for 16 hours. Haploid progeny were clonally isolated and genotyped at polymorphic markers across the genome (Supplemental Data). We show here genotyping results for four representative loci along supercontig 7 (sc7). All clones isolated from EPCM-treated cultures contained unrecombined parental genotypes, while clones isolated from VFCM-treated cultures showed clear evidence of ination. Top numbers show marker genomic positions along sc7.

Although swarming has not previously been reported in choanoflagellates and the biological significance of swarming in *S. rosetta* was not immediately obvious, swarming is associated with mating in diverse motile eukaryotes, including amoebae, ciliates, crustaceans, insects, fish, birds, and bats (Avery, 1984; Buskey, 1998; Downes, 1969; Giese, 1959; O'Day, 1979; Veith et al., 2004; Watson et al., 2003). Therefore, we hypothesized that swarming in *S. rosetta* might indicate mating. To investigate whether *V. fischeri*-induced swarming is a prelude to mating, we sought to determine whether the hallmarks of mating in microbial eukaryotes (cell fusion, nuclear fusion, and meiotic recombination (Bell, 1988; Dini and Nyberg, 1993; Levin and King, 2013)) occur in *S. rosetta* following treatment with *V. fischeri*.

Our lab previously found that starved *S. rosetta* cells mate at low frequencies (<2% of the population) after starvation for 11 days (Levin and King, 2013; Levin et al., 2014). In contrast, as early as 30 minutes after induction with live *V. fischeri* or conditioned medium isolated from a *V. fischeri* culture, *S. rosetta* cells formed swarms and then began to pair up and fuse (Figure 1C and Supplemental Movie 2). Once paired, cell fusion took as little as three minutes and all observed cell pairs were oriented with their basal poles (opposite the flagellum) touching. Paired cells subsequently fused along the basal pole, resulting in the formation of a binucleate cell harboring two flagella (Figure 1D). After cell fusion, the two nuclei congressed and fused, and one of the two parental flagella eventually retracted (Figure S1B), resulting in a diploid cell.

While cell fusion and nuclear fusion are consistent with mating, parasexual processes can occur in the absence of meiotic recombination (Goodenough and Heitman, 2014). Therefore, to test whether swarming was associated with the initiation of a true sexual cycle, we used *V. fischeri* to induce the formation of heterozygous diploids in *S. rosetta* cultures and then examined their offspring for evidence of meiosis and recombination (Figure 1E). To produce heterozygous diploid cells, two haploid *S. rosetta* strains (R+ and R–) containing previously characterized polymorphisms (Levin et al., 2014) were mixed either in the presence of *V. fischeri* conditioned medium (VFCM), or in conditioned medium from *Echinicola pacifica* (EPCM), a prey bacterium (Levin et al., 2014) that does not induce swarming, as a negative control (Figure 1E). After VFCM or EPCM exposure, 48 clones were isolated from each culture condition. Although we cannot directly measure the ploidy of live *S. rosetta* cells, heterozygous diploids can be readily identified by genotyping. While all clones (48/48) reared from the EPCM-treated culture contained un-recombined parental genotypes, 10/48 clones isolated following VFCM treatment were shown by genotyping to be heterozygous diploids. We surmised that the heterozygous diploids were the result of outcrossed mating, and found that further passaging of these clones yielded motile haploid progeny. 147 haploid progeny from three different heterozygous diploids were clonally isolated and genotyped at polymorphic sites across the genome, providing evidence for independent assortment and meiotic recombination (Figure 1E and Supplemental File 1). Taken together, these results demonstrate that *V. fischeri* produces an aphrodisiac that induces swarming and mating in *S. rosetta*.

### Bioactivity-guided fractionation revealed that the *V. fischeri* aphrodisiac is a protein

Automated image analysis of *S. rosetta* cells swarming in response to VFCM provided the basis for a quantitative bioassay of mating induction (Figure 2A,B and Methods). As a baseline, we found that 30 minutes after treatment with VFCM, *S. rosetta* cells consistently formed swarms containing between 5-35 cells each, whereas cells did not form clusters in response to EPCM. Using this bioassay, we first tested whether *V. fischeri* cues involved in quorum sensing (e.g. homoserine lactones) (Lupp and Ruby, 2005; Lupp et al., 2003) or those required for its symbiosis with the squid *Euprymna scolopes* (e.g. lipopolysaccharide (LPS) and peptidoglycan (PGN)) (Koropatnick et al., 2004; Shibata et al., 2012) might contribute to swarming induction in *S. rosetta*. A set of five different *V. fischeri* mutant strains that are deficient in quorum sensing were all wild type for swarming induction (Lupp and Ruby, 2005; Lupp et al., 2003), as were seven mutants in polysaccharide export pathways required for symbiosis with *E. scolopes* (Shibata et al., 2012) (Supplemental Table 1). Moreover, treatment of *S. rosetta* with purified quorum sensing molecules (Supplemental Table 2) and *V. fischeri* outer membrane vesicles (OMVs) containing LPS and PGN (Beemelmanns et al., 2014) (Figure S2A) also failed to elicit mating, suggesting that the cue(s) required for *S. rosetta* mating induction likely differ from factors required either for quorum sensing or squid colonization.

**Figure 2.**
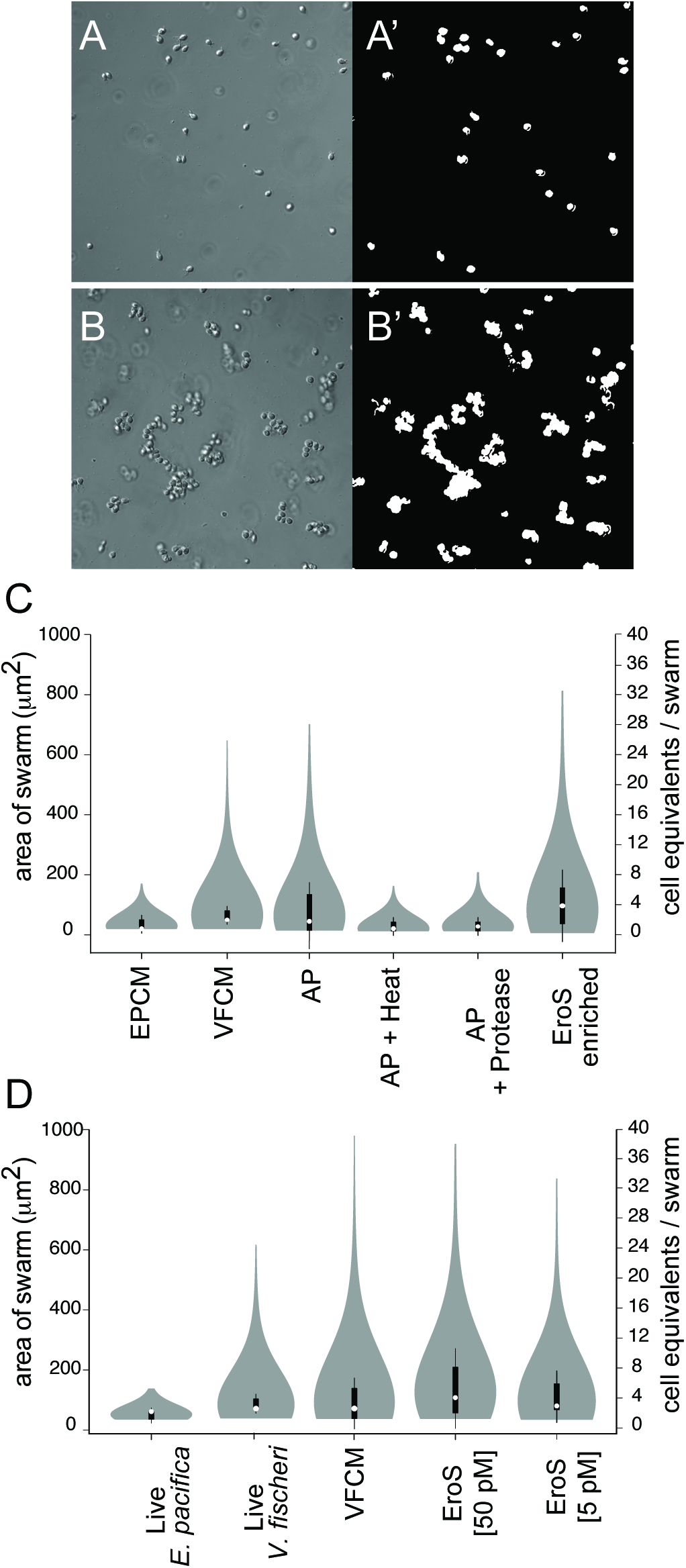
Bioactivity-guided isolation of the *V. fischeri* aphrodisiac. **(A, B)** Automated image analysis allowed quantification of *S. rosetta* swarming in response to *V. fischeri*-derived activity. Pictured are *S. rosetta* cells 30 minutes after treatment with *E. pacifica* conditioned media (A) or *V. fischeri* conditioned media (B). By generating a binary mask (A’, B’) we could measure the area of the swarm, and estimate the number of cells (“cell equivilants”) per swarm. **(C)** Swarming in *S. rosetta* is induced by compounds in the ammonium sulfate precipitation of *V. fischeri* culture supernatant (AP), but not by AP exposed to heat (80°C for 10 minutes; AP + Heat) or proteases (AP + Protease). The aphrodisiac activity tracked with a ∼90kD protein band that was revealed by mass spectrometry to be the *V. fischeri* EroS protein (VF_A0994). **(D)** EroS triggers mating at plausible environmental concentrations. Purified EroS induces swarming in *S. rosetta* at concentrations as low as 5 pM, and is sufficient to fully recapitulate the aphrodisiac activity of live *V. fischeri* bacteria and VFCM.

We next turned to an unbiased, activity-guided fractionation and found that the aphrodisiac was enriched in VFCM, including after depletion of OMVs (Figure S2A). The aphrodisiac produced by *V. fischeri* could be recovered from VFCM by ammonium sulfate precipitation, and the activity of the ammonium sulfate fraction was sensitive to both heat and protease treatment, suggesting that the activity might be proteinaceous (Figure 2C). We therefore separated all proteins precipitated from VFCM by size exclusion and anion exchange chromatography, and tested the protein fractions in the swarming bioassay (Figure 2A-C and Figure S2B). The swarming activity tracked with a single ∼90kD protein, which was revealed by mass spectrometry to be the uncharacterized *V. fischeri* protein VF_A0994, hereafter referred to as EroS for <u>E</u>xtracellular <u>r</u>egulator <u>o</u>f <u>S</u>ex (Figure 2C and Supplemental File 2; GenPept Accession YP_206952). To test whether EroS was sufficient to induce swarming in *S. rosetta*, we heterologously expressed the *eroS* gene in *E. coli* and found that the purified protein recapitulated the swarm-inducing activity of live *V. fischeri* and VFCM (Figure 2D). Purified EroS protein was also sufficient to induce mating between two *S. rosetta* strains (R+ and R–; Supplemental Table 3), demonstrating that a single protein secreted by *V. fischeri* is sufficient to induce both swarming and mating in *S. rosetta*.

### EroS protein is a chondroitinase

To understand the mechanism by which *V. fischeri* induces choanoflagellate mating, we set out to determine the biochemical function of EroS. The EroS protein sequence contains a predicted glycosaminoglycan (GAG) lyase domain (supported by the detection of PFAM domains PF08124, PF02278, PF02884; (Finn et al., 2016)). GAGs are linear polysaccharides that are integral components of the animal extracellular matrix (ECM). GAG lyases depolymerize GAGs through an elimination mechanism that distinguishes them from hydrolases, and are produced by a subset of primarily pathogenic bacteria and fungi (Zhang et al., 2010), as well as by human commensals, including gut bacteria in the genus *Bacteroides* (Ahn et al., 2011; Hong et al., 2002). Through the alignment of the EroS protein sequence with multiple bacterial GAG lyases with solved structures, we found that EroS harbors conserved residues at sites required for catalytic activity (His-278 and Tyr-287; Figure 3A) (Han et al., 2014; Linhardt et al., 2006; Shaya et al., 2008; Weijun Huang et al., 2001).

**Figure 3.**
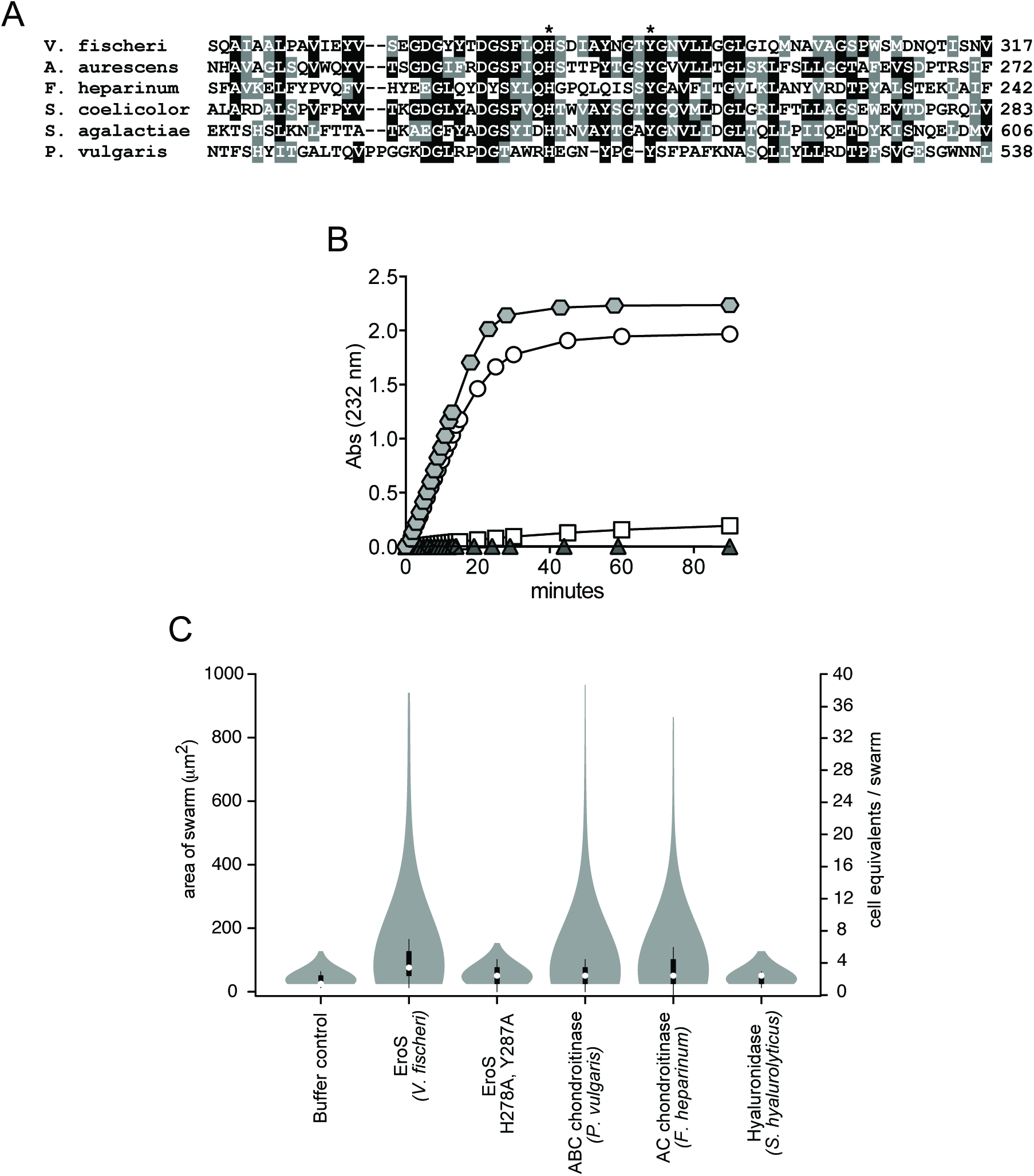
The *V. fischeri* aphrodisiac is a chondroitinase. **(A)** Alignment of the *V. fischeri* EroS amino acid sequence to diverse bacterial GAG lyases reveals that *V. fischeri* harbors conserved His and Tyr residues (indicated by *) at sites required for catalytic activity in characterized GAG lyases. Amino acids with >50% conservation between sequences are shaded (black shading for identical amino acids and grey shading for similar amino acids. **(B)** Purified EroS degrades chondroitin sulfate and hyaluronan. EroS was incubated with purified chondroitin sulfate (open circle), hyaluronan (grey hexagon), dermatan sulfate (open square), and heparan sulfate (grey triangle), and lyase activity of EroS was measured by monitoring the abundance of unsaturated oligosaccharide products with an absorbance at 232nm. Chondroitin sulfate and hyaluronan oligosaccharides accumulated rapidly in the presence of EroS, indicating depolymerization, whereas heparan sulfate and dermatan sulfate were not depolymerized by EroS. **(C**) The chondroitinase activity of EroS is necessary and sufficient for its function as an aphrodisiac. EroS protein with mutations in predicted catalytic resides (H278A, Y287A) fail to induce swarming in *S. rosetta. P. vulgaris* ABC chondroitinase and *F. heparinum* AC chondroitinase are sufficient to induce swarming at levels similar to EroS, whereas *S. hyalurolyticus* hyaluronidase fails to induce swarming, indicating that chondroitinase activity is necessary and sufficient for aphrodisiac activity.

Sulfated GAGs are thought to be eumetazoan-specific innovations (DeAngelis, 2002a; Yamada et al., 2011), and are not known to exist in choanoflagellates. (Although some pathogenic bacteria evade host immune responses by producing extracellular GAGs, these molecules are produced by way of an independently evolved biosynthetic pathway and, unlike animal GAGs, are not modified by sulfation (DeAngelis, 2002b)). Moreover, GAGs are diverse and the substrate specificities of GAG lyases cannot be deduced from sequence alone (Zhang et al., 2010). Therefore, we next set out to answer three questions: (1) does EroS exhibit GAG-degrading activity, (2) is the enzymatic activity of EroS required for its function as an aphrodisiac, and (3) what are its substrates in *S. rosetta*?

We found that purified EroS protein degrades GAG substrates *in vitro*, and is thus a *bona fide* GAG lyase. GAGs are classified based on their disaccharide units: heparan sulfate, chondroitin sulfate, dermatan sulfate, hyaluronic acid, and keratan sulfate (Zhang et al., 2010). EroS showed strong lyase activity toward purified chondroitin sulfate and hyaluronan, but not heparan sulfate or dermatan sulfate (Figure 3B and Figure S2C,D). We did not test keratan sulfate because it does not contain uronic acid and therefore cannot be degraded by GAG lyases (Garron and Cygler, 2010).

We next asked whether the enzymatic activity of EroS is important to its function as an aphrodisiac. Alanine substitution at conserved catalytic residues (His-278 and Tyr-287) required for chondroitin and hyaluronan degradation in other chondroitin lyases eliminated the ability of EroS to induce swarming in *S. rosetta* (Fig. 3C) (Linhardt et al., 2006). Moreover, well-characterized chondroitin lyases from other bacteria (ABC chondroitinase from *Proteus vulgaris* and AC chondroitinase from *Flavobacterium heparinum*) induced swarming and mating in *S. rosetta* at levels resembling those induced by EroS (Figure 3C and Supplemental Table 3), indicating that the chondroitinase activity of EroS is both necessary and sufficient for its function as an aphrodisiac.

Although sulfated GAGs were previously thought to be restricted to animals, key heparan biosynthetic enzymes have been detected in the genome of the choanoflagellate *Monosiga brevicollis* (Ori et al., 2011), and we have further found that the *S. rosetta* genome encodes homologs of enzymes required for chondroitin biosynthesis (Figure 4A, Figure S3A, Supplemental File 3) (Fairclough et al., 2013; King et al., 2008). To test whether chondroitin is produced by *S. rosetta*, we treated polysaccharides isolated from motile *S. rosetta* cells with the broad specificity ABC chondroitinase from *P. vulgaris*. The *P. vulgaris* ABC chondroitinase liberated chondroitin disaccharides, demonstrating that *S. rosetta* indeed produces chondroitin (Figure 4B, Figure S3B,C). Finally, to test whether EroS can degrade *S. rosetta* chondroitin we treated *S. rosetta* polysaccharides with EroS and found that it released unsulfated chondroitin and chondroitin-6-sulfate disaccharides, indicating that *S. rosetta* chondroitin is a target of EroS (Figure 4B, Figure S3B,C).

**Figure 4.**
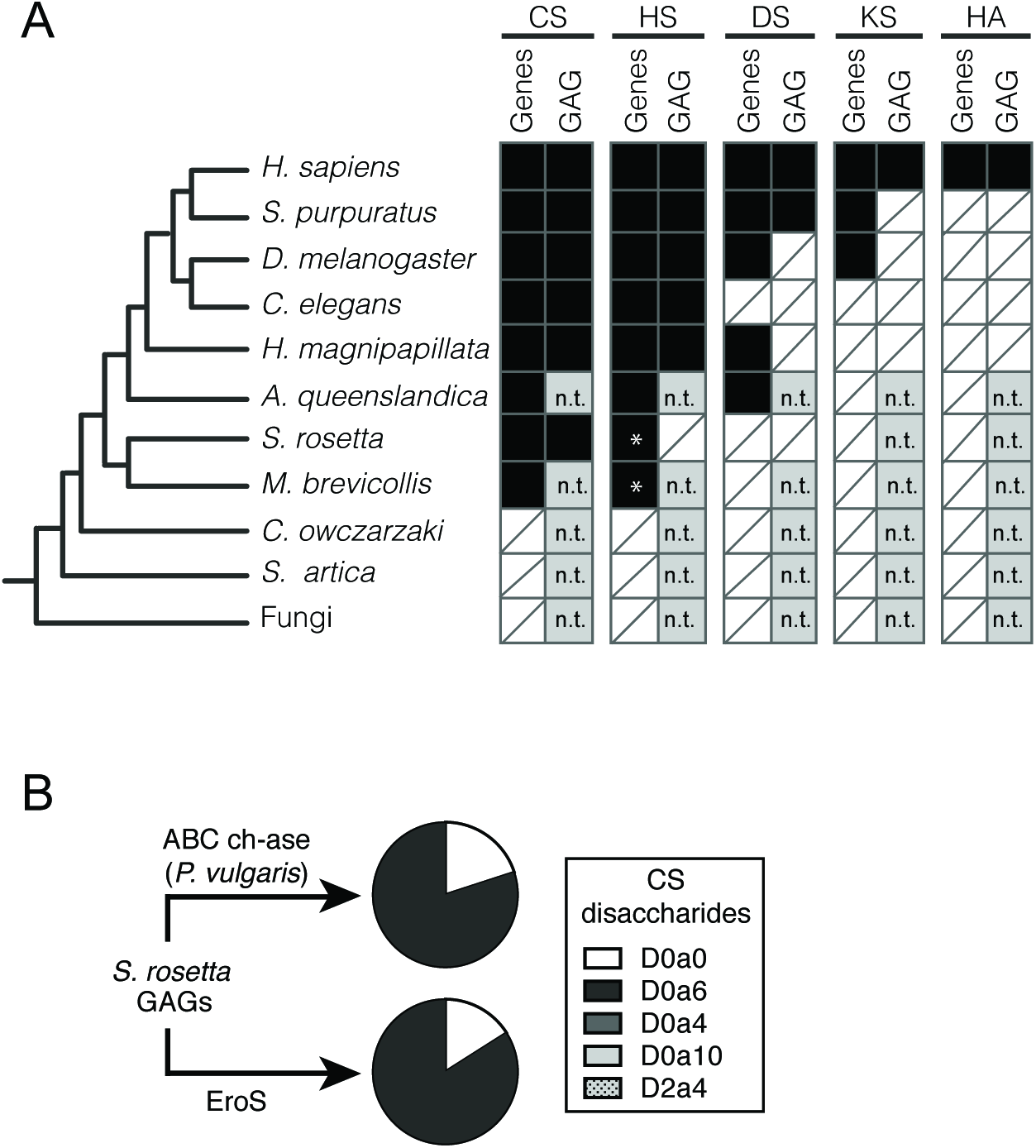
*S. rosetta* produces chondroitin sulfate that can be degraded by EroS. **(A)** Phylogenetic distribution of diverse GAGs [CS= chondroitin sulfate; HS= heparan sulfate; DS= dermatan sulfate; KS= keratan sulfate; HA= hyaluronan], and their biosynthetic genes. The presence (black box) and absence (white box with slash) of genes required for the biosynthesis of GAGs (Gene) and biochemical evidence for GAGs (Expt) in select opisthokonts. *; Ori et al. (2011) identified putative homologs of a subset of HS biosynthetic enzymes in the *M. brevicollis* genome, and we detect homologs of the same limited set of HS biosynthetic enzymes in *S. rosetta*. Importantly, these enzymes are shared components of the chondroitin biosynthetic pathway, and digestion of *S. rosetta* polysaccharides with heparinases failed to liberate heparan sulfate disaccharides, suggesting *S. rosetta* does not produce heparan sulfate (also refer to Figure S3). n.t.; not tested (experiments have not been performed to biochemically profile GAGs). **(B)** *S. rosetta* produces chondroitin that can be degraded by EroS. Polysaccharides isolated from *S. rosetta* were treated with either *P. vulgaris* ABC chondroitinase, an enzyme that can degrade many modifications of chondroitin into its disaccharide units (CS disaccharides), or purified EroS. Both ABC chondroitinase and EroS yielded similar amounts of unsulfated chondroitin disaccharide (D0a0) and chondroitin-6-sulfate disaccharide (D0a6) degradation products, indicating that unsulfated chondroitin and chondroitin-6-sulfate are produced by *S. rosetta*. In contrast, we were unable to detect chondroitin-4-sulfate (D0a4), chondroitin-4,6-sulfate (D0a10), or chondroitin-2,4-sulfate (D2a4) following degradation of *S. rosetta* polysaccharides with either EroS or ABC chondroitinase.

Chondroitin sulfate in animal ECM can be found covalently linked to core proteins, thus forming proteoglycans. To determine whether chondroitin disaccharides released from proteoglycans play a role in stimulating mating, we tested various products of EroS digestion for aphrodisiac activity (Figure S4). Conditioned media from EroS-treated *S. rosetta* cells did not trigger swarming in naïve *S. rosetta*, nor did the digested products of commercial chondroitin sulfate treated with EroS. Moreover, swarming was not induced by any combination or concentration of unsulfated and 6-sulfated chondroitin disaccharides tested (Figure S4). These results lead us to hypothesize that the structural modification of *S. rosetta* proteoglycans by EroS, rather than the chondroitin disaccharide products of EroS digestion, are important for activating the swarming and mating pathway in *S. rosetta*.

Finally, because swarming has not been previously described in choanoflagellates, we investigated whether *V. fischeri* might induce swarming under plausible environmental conditions. We found that EroS is secreted constitutively by *V. fischeri* when grown under either high or low nutrient conditions (Supplemental Table 4). Cultures of *S. rosetta* swarm in response to as little as 4×10^2^ *V. fischeri* cells/mL – a density comparable to that of *V. fischeri* in oligotrophic oceans (from 6×10^2^ cells/mL to >1×10^4^ cells/mL during blooms (Jones et al., 2007)) – within 30 minutes of exposure (Figure S5). Moreover, EroS was sufficient to trigger robust swarming in *S. rosetta* at concentrations as low as 5 pM (Figure S5), making EroS as potent as the sex pheromones produced by volvocine algae (Kochert, 2012) and by marine invertebrates (Bartels-Hardege et al., 1996; Li et al., 2002). Together, these data suggest that *V. fischeri* or other chondroitinase-producing bacteria could plausibly trigger *S. rosetta* swarming and mating *in natura*.

## Discussion

We have discovered that a secreted bacterial chondroitinase, EroS, induces mating in *S. rosetta*, one of the closest living relatives of animals. Through the study of this unexpected interkingdom interaction, we found that mating in *S. rosetta* is initiated in response to the degradation of chondroitin sulfate, a glycosaminoglycan previously thought to be restricted to animals.

The first hint that *V. fischeri* might induce mating came from the observation of *S. rosetta* swarms following exposure to the bacterium. By increasing local population density, swarming has previously been found to facilitate mating in diverse amoebae, flagellates, crustaceans, cnidarians, polychaetes, insects, fish, and birds (Avery, 1984; Buskey, 1998; Downes, 1969; Giese, 1959; Hamner and Dawson, 2009; Omori and Hamner, 1982; Sorensen and Wisenden, 2015; Watson et al., 2003). As in other organisms that swarm, the connection of swarming to mating may be critical, since their aquatic, pelagic lifestyle can make it challenging to find mates. Indeed, under the starvation conditions that trigger *S. rosetta* mating in the absence of swarming, mating takes >500X longer (∼11 days) and occurs in a small fraction of the population (Levin et al., 2014).

Most previously characterized examples of coordinated mating behaviors are regulated by pheromonal cues. Conspecific swarming is initiated by diverse aggregation pheromones (for example, ester and isoprenoid pheromones in beetles (Kartika et al., 2015; Wertheim et al., 2005) and peptide pheromones in sea slugs (Painter et al., 2016) and polychaetes (Ram et al., 1999)), and free-spawning marine animals produce pheromones to synchronize gamete release and enhance fertilization success (Babcock et al., 2011). Biotic and abiotic cues from the environment can also help coordinate mating behavior in animals. For example, spawning in marine invertebrates is correlated with phytoplankton blooms, and some sea urchins, mussels, and polychaetes spawn after exposure to small molecules produced by environmental phytoplankton (Smith and Strehlow, 1983; Starr et al., 1990; 1992), although these cues remain structurally elusive. Just as phytoplankton blooms are hypothesized to signify a nutrient-rich environment for spawning, the presence of chondroitinase-producing bacteria may indicate an environmental condition, or the convergence of multiple environmental factors, that favor mating in *S. rosetta*. Although *V. fischeri* was the first bacterium observed to regulate mating in *S. rosetta*, we have since identified other bacteria that similarly induce swarming and mating (Supplemental Table 1). Therefore, we predict that mating in *S. rosetta* might be regulated by diverse species of bacteria in nature, and hypothesize that swarming is a common occurrence within the natural life history of *S. rosetta*.

Our discovery that *V. fischeri* produces a chondroitinase that functions as an aphrodisiac also revealed that *S. rosetta* produces chondroitin sulfate, providing the first biochemical evidence for this important GAG in a non-animal and extending its evolutionary history to the premetazoan era. In an interesting parallel to the induction of *S. rosetta* mating by a chondroitinase, GAGs and sulfated polysaccharides mediate mating in diverse internally and externally fertilizing animals where they provide a protective and species-specific coating around oocytes (Miller and Ax, 1990). In the case of the mammalian oocyte, which is surrounded by the GAG hyaluronan, sperm secrete hyaluronidase to penetrate the hyaluronan-containing coating, whereas in sea urchins, sulfated polysaccharides coating the sea urchin oocyte ensure species-restricted sperm activation and binding (Mengerink and Vacquier, 2001). Of course, GAGs like hyaluronan and chondroitin sulfate are also essential components of the ECM in somatic cells of animals, where they contribute to a range of functions that include the maintenance of cell adhesion through interactions with ECM molecules, the integration of signals from the extracellular milieu, and the stabilization of collagen fibers. Future investigations may reveal whether chondroitin sulfate in *S. rosetta* functions to mediate species-specific cell recognition in the context of fertilization and may provide insight into the premetazoan roles of this important molecule. Moreover, it will be fascinating to explore whether bacteria influence mating in other aquatic organisms, for whom the triggers of mating are often obscure.

## Acknowledgements

We thank K. Visick for *V. fischeri syp* deletion mutants, N. Ruby for *V. fischeri* quorum sensing deletion mutants, and Y. Boucher for diverse species of *Vibrio* bacteria. We thank M. Abedin, D. Booth and B. Larson for helpful discussions and critical reading of the manuscript. This work used the Vincent J. Coates Proteomics/Mass Spectrometry Laboratory at UC Berkeley, supported in part by NIH S10 Instrumentation Grant S10RR025622, and the Complex Carbohydrate Research Center, supported by the National Institutes of Health-funded Research Resource for Integrated Glycotechnology (NIH grant no. P41GM103390).

## Methods

### Culture media

Artificial seawater (ASW), cereal grass media (CG media), and Sea Water Complete media (SWC) were prepared as described previously (Levin and King, 2013; Woznica et al., 2016). Artificial sea water (ASW) was made by adding 32.9 g Tropic Marin sea salts (Wartenberg, Germany) to 1L distilled water to a salinity of 32-27 parts per thousand. SWC media was made by adding 250 mg/L peptone, 150 mg/L yeast extract, 150 L/L glycerol in artificial sea water. CG media was made by infusing ASW with cereal grass pellets (Basic Science Supplies, Rochester NY).

### Choanoflagellate husbandry

SrEpac (Levin and King, 2013) (*S. rosetta* grown in the presence of *Echinicola pacifica* bacteria, ATCC PRA-390) was propagated in 5% Sea Water Complete media (SWC diluted to 5% vol/vol in ASW) at 22°C. SrEpac was passaged 1:20 into 19mL fresh 5% SWC every other day to obtain stationary growth phase cultures (cells were grown in 25cm^2^ Corning cell culture flask). Prior to all induction bioassays, unless otherwise indicated, cells were diluted to approximately 1×10^5^ cells/mL in ASW at the time of induction.

### Immunofluorescence microscopy

Stationary-phase cells were induced with *V. fischeri* conditioned media or *E. pacifica* conditioned media and fixed at intervals of 10 minutes, 30 minutes, 1 hour, 2 hours, and 4 hours after induction. After vortexing, cells were fixed for 5 min in 6% acetone followed by 10 minutes in 4% formaldehyde. Cells were then allowed to settle for 30 min onto poly-L-lysine coated coverslips (BD Bioscience). Cells were stained with E7 anti-tubulin antibody (1:200; Developmental Studies Hybridoma Bank), Alexa Fluor 488 anti-mouse (1:1000; Molecular Probes), and .01mg/mL Hoechst 3342 (Thermo Fischer) before mounting in Prolong Gold antifade reagent (Molecular Probes). Cells were imaged at 63x using a Zeiss LSM 880 AxioExaminer with Airyscan.

Mating stages were assigned based on the following criteria: orientation of paired cells, fusion of cell bodies, localization and number nuclei, number of flagella. Cell fusion could be clearly distinguished from cell division for several reasons, including (1) fusing cells are paired basally, whereas recently divided sister cells are paired laterally, (2) flagella remain elongated during the fusion process, but are retracted throughout cell division and (3) DNA remains uncondensed throughout cell fusion, but is condensed during cell division.

### Isolation of conditioned media (including VFCM and EPCM)

*Vibrio fischeri* ES114 (ATCC 700601) and all other *Vibrio* species (Supplemental Table 2) were grown by shaking in 200mL 100% SWC media for 30H at 20°C, and pelleted by centrifugation. *E. pacifica* was grown by shaking in 200mL 100% SWC for 30H at 30°C, and pelleted by centrifugation. Cell-free supernatant was then vacuum filtered twice through a 0.22 mM filter (EMD Millipore Stericup) to obtain 100% CM. Concentrated conditioned media was obtained using 30kD and 50kD molecular weight cut off centrifugal filter units (Amicon).

### Inducing mating and meiosis

#### *S. rosetta* strains

All crosses were performed between two *S. rosetta* strains with previously verified single nucleotide polymorphisms (SNPs), R-(previously referred to as Rosetteless) and R+ (previously referred to as Isolate B) (Levin et al., 2014). Prior to inducing mating, stationary phase cultures were obtained by passaging Rosetteless 1:20 into 19mLs fresh 5% SWC media every other day, and Isolate B 1:10 into 20mLs fresh 25% CG media every two days.

#### Inducing mating

Stationary phase R+ and R-cultures were counted and diluted to the same cell density (1×10^6^ cells/mL). R+ and R-cultures were mixed in equal proportions, pelleted, and resuspended in fresh 25% CG media to obtain a final cell density of 1×10^6^ cells/mL. Mating crosses were performed in 2mL total volumes under the following induction conditions: 5% (V/V) *E. pacifica* conditioned media (EPCM), 5% (V/V) *Vibrio fischeri* conditioned media (VFCM), 0.5nM VF_rGAG lyase, 5 “units” Chondroitinase ABC (Sigma C3667), and 5 “units” Chondroitinase AC (Sigma C2780). Cells were allowed to mate for 16 hours, after which the induced culture was pelleted and washed twice in 25% CG media to prevent further mating prior to limiting dilution.

#### Isolating diploids by limiting dilution

Mated cells were clonally isolated by limiting dilution into 96-well plates containing 25% CG media. For all crosses performed, the probability of clonal isolation at this step was between 0.85 and 0.92. Although we cannot directly measure the ploidy of live *S. rosetta* cells, the differentiation of planktonic motile cells into substrate-attached “thecate” cells correlates with the transition to diploidy (Levin et al., 2014). After five days of growth, isolates were phenotyped and then divided into two populations. For each isolate, one population was rapidly passaged to induce meiosis (see below), and the other population was used for DNA extraction. For DNA extraction, isolates were expanded into 1mL of 5% CG media to prevent meiosis, and grown for three days in 24-well plates. DNA was extracted from each isolate using the following method: 500µL of cells were pelleted and resuspened in 20µL base solution (25mM NaOH, 2mM EDTA). Base solutions from the isolates were transferred to a PCR plate, boiled at 100°C for 20 min, and cooled at 4°C for 5 min. 20µL Tris solution (40mM Tris-HCl, pH 7.5) was then added to each sample. 1µL of this sample was used as the DNA template for genotyping reactions. To identify which isolates were the result of outcrossed mating, isolates were genotyped at two unlinked microsatellite markers that are polymorphic between the R+ and R-parental strain (Levin et al., 2014). All outcrossed diploids isolated were phenotypically thecate as opposed to motile planktonic. No thecate isolates were observed in control EPCM treated cultures.

#### Isolation of haploid meiotic progeny

Immediately after phenotyping, clones isolated by limiting dilution were passaged 1:10 into 1mL fresh 25% CG media to induce meiosis. Thecate clones that were outcrossed diploids typically gave rise to a clear mixture of haploid chains and rosettes after two days. Haploids were clonally isolated by limiting dilution into 96-well plates containing 25% CG media, and phenotyped after five days. Meiosis was confirmed either by 1) genotyping at two unlinked microsatellite markers, or 2) by genotyping at 38 markers using KASP technology (LGC Genomics, Beverly, MA) (Levin et al., 2014).

#### Genotyping meiotic progeny

To confirm genome-wide recombination, haploid progeny isolated from the 5% VFCM-induced cross were genotyped at 38 markers (Levin et al., 2014). Briefly, three outcrossed diploids (named A2, A3, and H2) were rapidly passaged to induce meiosis, and clones from each outcrossed diploid were isolated by limiting dilution. The probability of clonal isolation at this step was .94 for A2, .93 for A3, and .91 for H2. A total of 147 haploid isolates from the three outcrossed diploids were phenotyped and expanded for subsequent DNA extraction and genotyping.

### Quantifying mating swarms

Inductions were set up in 100µL volumes in 96-well glass bottom plates (Ibidi 89626). Assays were imaged at 10X magnification using transmitted light (bright field) on the Zeiss Observer Z.1 platform using a Hamamatsu C11440 camera. An automated sequence was set up such that each sample was imaged at 4 distinct locations throughout the well.

Images were batch processed in ImageJ to ensure consistency. After applying the ‘Smooth’ command to reduce background bacterial signal, the ‘Find Edges’ command was applied to further highlight the phase-bright choanoflagellate cells. Images were then converted to black and white using the ‘Make Binary’ command, followed by the “Close” command to fill in small holes.

Finally, images were analyzed using the ‘Analyze Particles’ command to calculate the area of each cell or swarm (the white space in Figure 2A’,B’) within an image.

### Isolating the *Vibrio fischeri* mating induction factor

#### Preparation of >30kD-enriched VCM

Eight 1L cultures of *V. fischeri* ES114 were grown shaking for in 100% SWC for 24 h at 25°C. Cultures were pelleted at 16,000 × g, and the supernatants were concentrated to 120 mL using a tangential flow filtration device with a 30 kDa centramate filter (Pall #OS030T12). The supernatant was then further clarified by pelleting 39,000 × g.

#### Ammonium sulfate precipitation

>30kD-enriched VCM was treated with 1 M Tris-HCl (pH 7.6) and fractionally precipitated with increasing (40%-75%) concentrations of ammonium sulfate. Precipitates were resuspended in water, and tested in the swarming bioassay.

#### Size exclusion chromatography

Active ammonium sulfate precipitation fractions were combined and concentrated to 1 mL. 0.85 mL was injected on a HiPrep^TM^ 16/60 Sephacryl^TM^ S-200 High Resolution column (GE Healthcare Life Sciences #17-1166-01) using an AKTA Explorer FPLC instrument. Proteins were eluted with 30 mM Tris-HCl (pH 7.7, 4 °C) at 0.5 mL/min for 120 mL, and 2 mL fractions were collected. Adjacent fractions were paired and tested in the swarming bioassay, as well as analyzed by PAGE.

#### Anion exchange chromatography

Active SEC fractions were combined and concentrated to 1 mL in Solvent A (20 mM L-histidine, pH 6.0) and injected at 2.5 mL/min into a HiPrep^TM^ 16/10 Q XL column (GE Healthcare Life Sciences #17-5092-01). Proteins were eluted in 2 mL fractions over a 300 mL linear gradient (0-100%) of Solvent A to Solvent B (1 M NaCl in 20 mM L-histidine, pH 6.0). Fractions were tested in the swarming bioassay and analyzed by PAGE.

#### Renaturing proteins after PAGE

Proteins from highly active AEX fractions were concentrated and mixed with 15 µg/mL β-lactoglobulin carrier protein, and run in adjacent lanes through a NuPAGE^TM^ 4-12% Bis-Tris polyacrylamide gel. Evenly spaced bands were excised from one lane, and the remaining gel was stained with Coomassie blue R-250 and retained for mass spectrometry. The excised slices were crushed, and then extracted for 6 hours with 300 µL of elution buffer (50 mM Tris-HCl pH 7.7, 100 µM EDTA, 1 mM DDT, 150 mM NaCl, 0.1% sodium dodecyl sulfate [SDS], 100 µg/mL bovine serum albumin [BSA], pH 7.7) (Hager and Burgess, 1980). Proteins were precipitated with 1200 µL cold acetone, incubated on dry ice for 30 minutes, and pelleted at 16,000 × g for 10 minutes at 4 °C. Air-dried pellets were dissolved in 10 µL of solubilization buffer (50 mM Tris-HCl pH 7.7, 100 µM EDTA, 1 mM DDT, 150 mM NaCl, 20% glycerol, 6 M guanidine hydrochloride) for 20 minutes, and then diluted with 500 µL of dilution buffer (50 mM Tris-HCl pH 7.7, 100 µM EDTA, 1 mM DDT, 150 mM NaCl, 20% glycerol). Proteins further renatured for 3.5 H at room temperature, and were concentrated to 20 µL. Proteins excised from bands were then tested in the swarming bioassay.

#### Mass Spectrometry

Because only one excised protein band displayed bioactivity, the corresponding slice retained for mass spectrometry was subjected to trypsin digestion and LC-MS/MS at the Proteomics/Mass Spectrometry Laboratory at UC Berkeley. Complete mass spectrometry results are listed in Supplemental File 2.

### Protein expression and purification

The *eroS* gene was amplified by PCR (Phusion^®^ DNA polymerase, New England Biosciences) from *V. fischeri* ES114 genomic DNA (Forward primer: 5’-GCCTCTGTCGACGCAAAAAATACCCAAACACCAC; Reverse primer: 5’-AATTAAGCGGCCGCCGTCTTGAATTGTTACTTGGAAAGAATAAG). After digestion with *SalI-*HF and *NotI*-HF (New England Biolabs), the *eroS* gene was ligated in-frame into a pET6xHN-N vector (Clontech) for fusion to a His-tag and transformed into OneShot BL21(DE3) cells (Invitrogen) for expression. Transformed *E. coli* were grown at 37 °C, 200 rpm shaking in LB media supplemented with 100 µg/mL ampicillin. After growth to OD 1.0, the temperature was decreased to 16 °C, and protein expression was induced by addition of 1 mM IPTG. After 24 hours, cells were pelleted and lysed with xTractor^TM^ buffer (Clontech), purified with HisTALON^TM^ gravity columns (Clontech), and the His-tag was released by enterokinase cleavage (Millipore enterokinase cleave capture kit #69067).

A gBlock of the *eroS* gene sequence harboring H278A and Y287A mutations was purchased from Integrated DNA Technologies and ligated into the pET15b vector (Novagen) at *NdeI* and *BamHI* restriction sites. The mutant *eroS* gene was transformed into OneShot BL21 Star (DE3) cells (Invitrogen) for expression. Transformed *E. coli* were grown at 37 °C, 220 rpm shaking in LB media supplemented with 100 µg/mL ampicillin. After growth to OD 0.8, the temperature was decreased to 16 °C, and protein expression was induced by addition of 0.3 mM IPTG. After 24 hours, cells were lysed and protein was purified with HisPur Cobalt Resin (Thermo Scientific).

### Amino acid sequence alignments

Amino acid sequences from *V. fischeri* (VF_A0994, GenBank: AAW88064.1) and characterized bacterial GAG lyases [*A. aurescens* (AC lyase, PDB: 1RWG_A), *S. coelicolor* (AC lyase, PDB: 2WDA_A), *S. agalactiae* (hyaluronate lyase, PDB: 1LXM_A), *F. heparinum* (AC lyase, PDB: 1CB8_A), and *P. vulgaris* (ABC chondroitinase, GenBank: ALL74069.1)] were aligned using Clustal Omega multiple sequence alignment (http://www.ebi.ac.uk/Tools/msa/clustalo/). Conserved amino acid residues were highlighted using BoxShade (http://www.ch.embnet.org/software/BOX_form.html).

### EroS GAG lyase activity *in vitro*

The glycosaminoglycan cleavage activity of purified Eros was determined *in vitro* as previously described (Wang et al., 2015). Briefly, GAG standards [hyaluronic acid (Sigma #H5388), chondroitin sulfate (Sigma #C4384), dermatan sulfate (Sigma #C3788), and heparin (Sigma #H3393)] were dissolved to a concentration of 1 mg/mL in buffer solution (50 mM NaH_2_PO_4_/Na_2_HPO_4_, 0.5 M NaCl, pH 8.0). 1 mL of GAG standard was added to 5 µL of enzyme solution. The assays were performed in quartz cuvettes (1 cm pathlength) at 23 °C, and UV absorbance measurements (232 nm) were taken directly on the reacting mixture.

### AQUA quantification of EroS

Absolute Quantification (AQUA) peptide (Gerber et al., 2003) was used to accurately quantify the concentration of purified EroS and EroS in *Vibrio fischeri* conditioned media. Briefly, *V. fischeri* was grown for 8 H in 100% SWC media and 5% SWC media. Purified EroS and concentrated *Vibrio fischeri* culture supernatants were loaded onto a PAGE gel. The gel was stained with Coomassie blue R-250, and bands containing VF_A0994 were excised and sent to the Taplin Mass Spectrometry facility (Harvard Medical School) for further analysis. Synthetic AQUA peptide (TQITDDTYQNFFD[KC13N15], Sigma-Aldrich) and trypsin were added to each excised band, and LC-MS/MS was performed on the digested peptides using a Thermo Scientific Orbitrap. The amount of EroS present in each gel slice was calculated by comparing MS2 peak intensities of the native peptide with the internal AQUA synthetic peptide standard.

### *S. rosetta* polysaccharide isolation and GAG disaccharide analysis

6×500 mL cultures of SrEpac were grown in 5% SWC until mid-stationary phase, and washed 3x to reduce bacterial load before being pelleted, flash frozen, and lyophilized. 125 mg of lyophilized *S. rosetta* sample was sent to the Complex Carbohydrate Research Center for GAG isolation, digestion, and SAX-HPLC.

Polysaccharides were isolated from the *S. rosetta* sample and digested with either EroS or chondroitinase ABC (Sigma C3667) for chondroitin disaccharide analysis, and heparinases I, II, and III (Dextra Laboratories) for heparan disaccharide analysis. Briefly, a ratio of 10 µL *S. rosetta* polysaccahrides to 1uL of enzyme was incubated for 24 hours. Samples were heated to 100°C for 5 minutes to inactivate the enzyme, and centrifuged at 14,000 rpm for 30 minutes prior to SAX-HPLC.

SAX-HPLC was carried out on an Agilent system using a 4.6×250 mm Waters Spherisorb analytical column with 5µm particle size at 25°C. Detection was performed by post-column derivatization. Briefly, the eluent from the column was combined with a 1:1 mixture of 0.25 M NaOH and 1% 2-cyanoacetamide pumped at a flow rate of 0.5 mL/min from a post-column reactor. The eluent was heated to 130°C in a 10m reaction coil, then cooled in a 50-cm cooling coil and directed into a Shimadzu fluorescence detector (λex=346 nm, λem=410). Commercial standard disaccharides (Dextra Laboratories) were used for identification of each disaccharide based on elution time, as well as calibration.

### Testing bioactivity of chondroitin disaccharides

Chondroitin disaccharides and chondroitin sulfate were tested for bioactivity by treating *S. rosetta* with unsulfated chondroitin disaccharides, chondroitin-6-sulfate disaccharides, unsulfated chondroitin + chondroitin-6-sulfate disaccharides, and chondroitin sulfate (from shark cartilage) at concentrations ranging from 0.0001M-0.1M. Cells were imaged and quantified after 30 minutes, 1 hour, and 3 hours.

Degradation products of chondroitin sulfate were generated by incubating 100 µg of chondroitin sulfate with 1µL of either EroS or ABC chondroitinase (*P. vulgaris*) overnight. Enzymatic activity was killed by incubating samples at 80°C for 5 minutes. The resulting degradation products were tested for bioactivity at concentrations ranging from 0.0001M-0.1M. Cells were imaged and quantified after 30 minutes, 1 hour, and 3 hours.

## Supplemental Information

**Figure S1.**
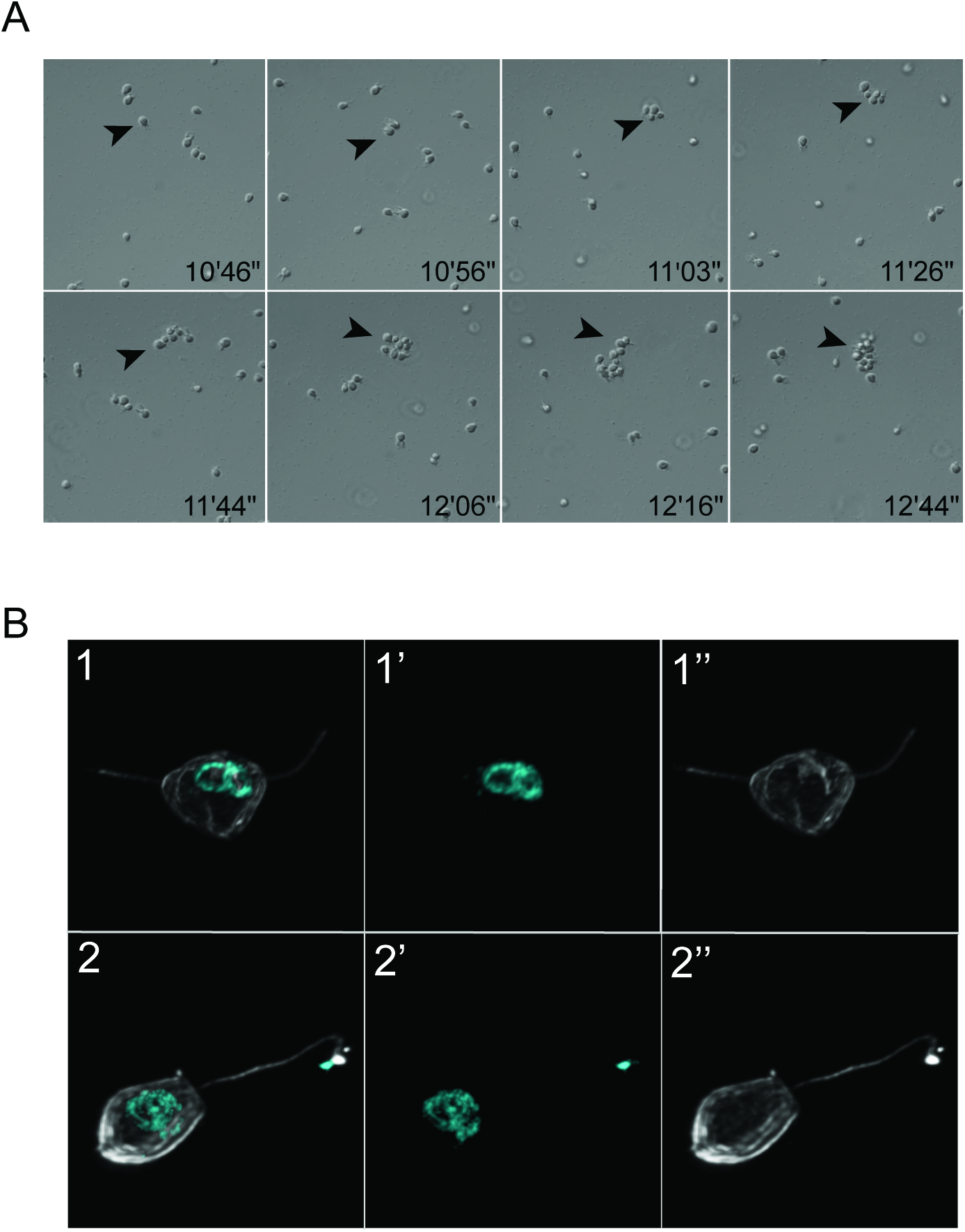
(related to Figure 1). *V. fischeri* induces swarming and mating in *S. rosetta*. **(A)** Stills of swarm formation after induction with *V. fischeri* bacteria. Arrowhead tracks the formation and movement of a single swarm over time. **(B)** Nuclear fusion in mating pairs of *S. rosetta* following treatment with *V. fischeri*. Pictured are late stages of nuclear congression and fusion. Following cell fusion, the nuclei congress towards the center of the bi-flagellated cell (B1-1’’), and fuse (B2-2’’). The final result of nuclear fusion is a diploid cell, harboring a single flagellum (B2-2’’). Hoechst (B1’,2’; cyan) highlights the nucleus, and anti-tubulin antibody (B1’’,2’’; white) highlights the cell body and flagellum.

**Figure S2.**
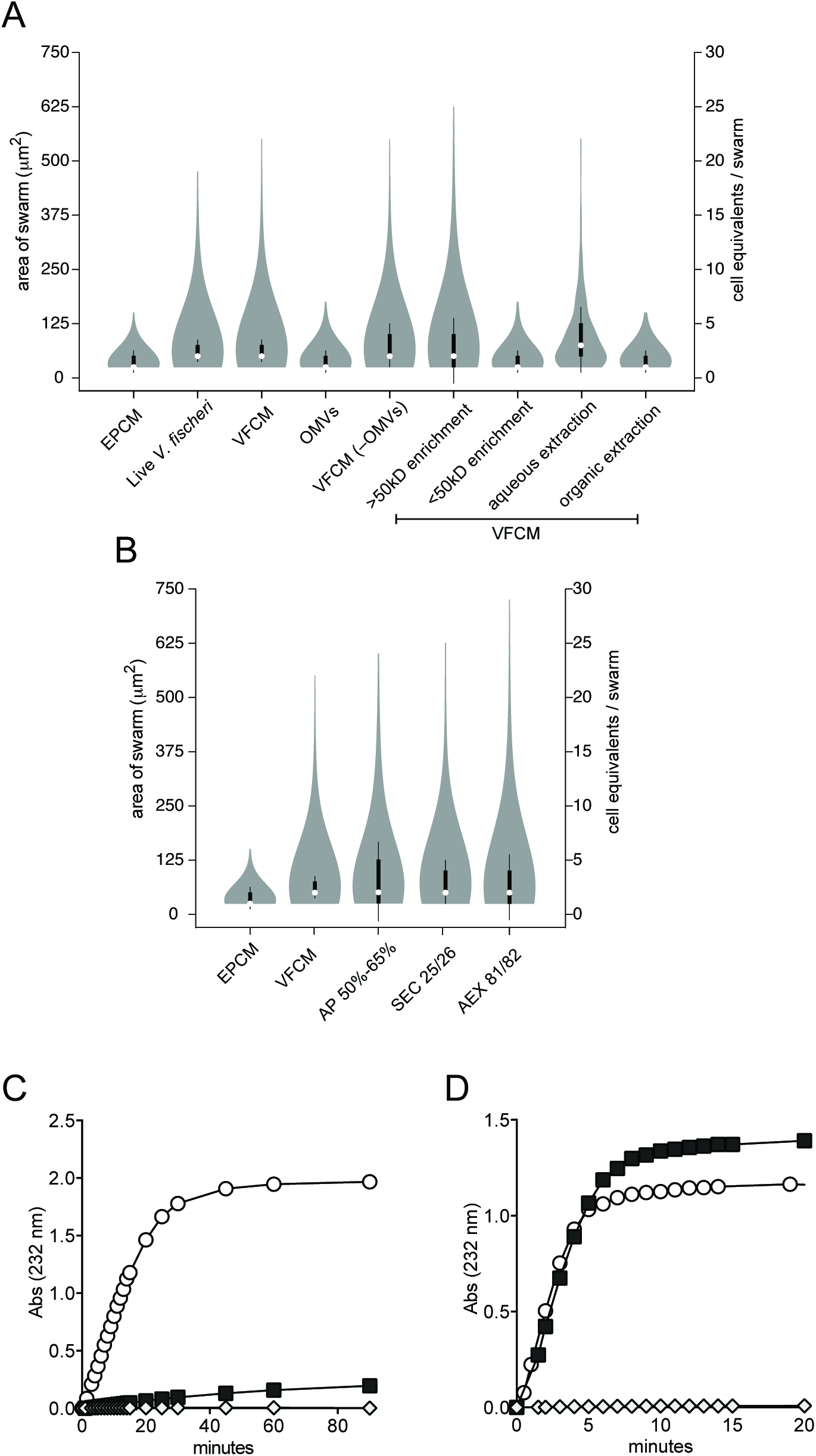
(related to Figure 2, Figure 3). Bioactivity-guided isolation and characterization of EroS. **(A)** Swarming in *S. rosetta* is induced by large (>50kD), water-soluble factors present in *V. fischeri* conditioned media (VFCM). **(B)** Isolation of EroS from VFCM. To identify the source of the aphrodisiac activity, proteins were precipitated from VFCM (AP 50%-65%) and separated by size exclusion (SEC) and anion exchange (AEX) chromatography. A protein band of ~90kD, later determined to be EroS (VF_A0994), was abundant in the bioactive SEC (SEC 25/26) and AEX (AEX 81/82) fractions. **(C, D)** EroS is a chondroitin AC lyase. **(C)** EroS degrades chondroitin sulfate AC, but not chondroitin sulfate B (dermatan sulfate) *in vitro*. EroS was incubated with purified chondroitin sulfate AC (open circle) and chondroitin sulfate B (dark grey square), and lyase activity of EroS was measured by monitoring the abundance of unsaturated oligosaccharide products with an absorbance at 232nm. Diamond represents a no enzyme control. **(B)** Chondroitinase ABC (*P. vulgaris*), a positive control for *in vitro* chondroitin degradation assays, rapidly depolymerizes both chondroitin sulfate AC (open circle) as well as chondroitin sulfate B (grey square). Diamond represents a no enzyme control.

**Figure S3.**
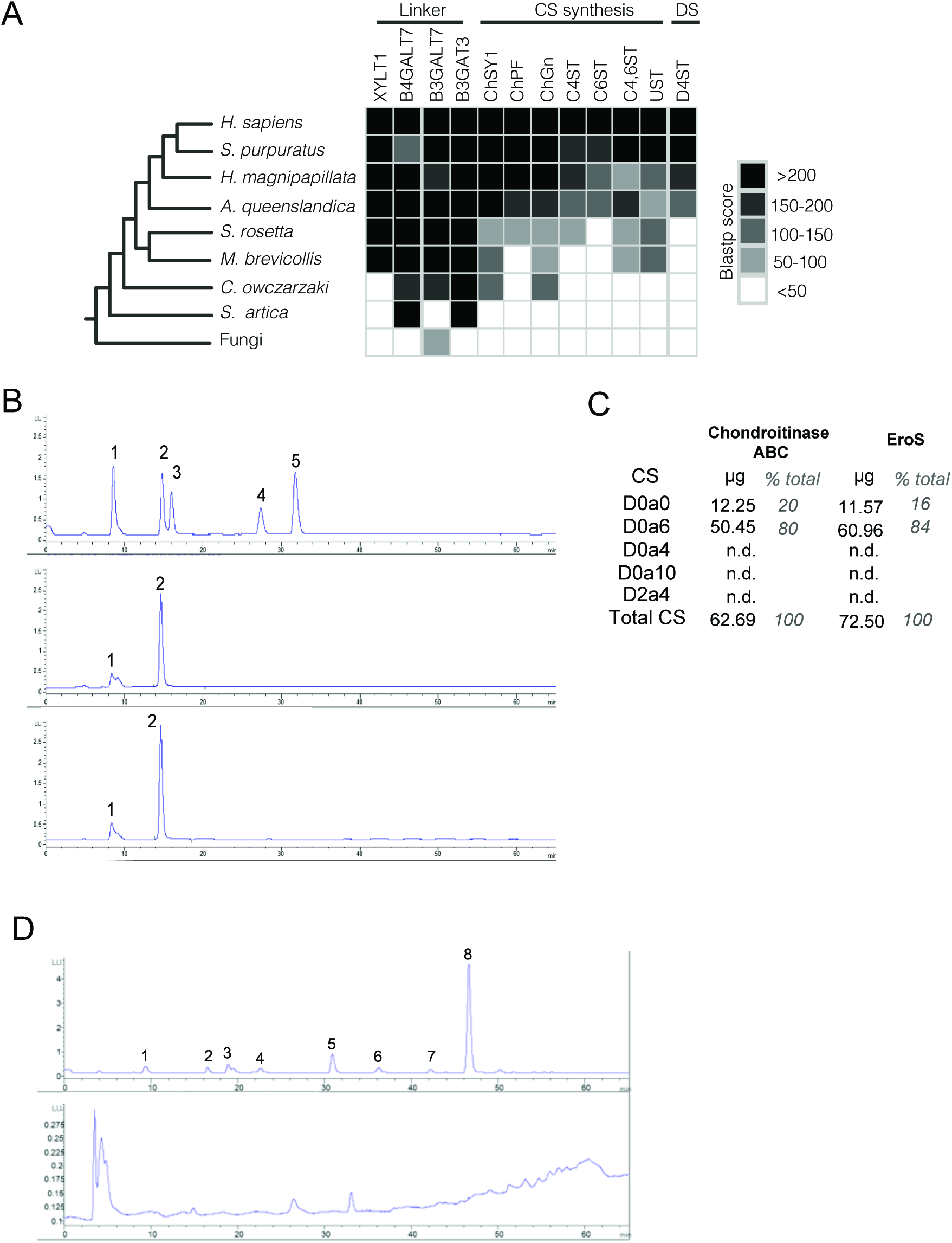
Chondroitin sulfate produced by *S. rosetta* can be degraded by EroS. **(A)** Orthologs of chondroitin sulfate (CS) synthesis, but not dermatan sulfate (DS) synthesis, are present in the genomes of choanoflagellates *S. rosetta* and *M. brevicollis*. Genes identified as “linker” synthesize the proteoglycan linker tetrasaccharide and are important for the biosynthesis of multiple types of GAG, whereas the genes identified as “CS synthesis” are specific to CS biosynthesis. The gene identified as “DS” is required for the formation of dermatan sulfate. All query sequences used were human orthologs. If multiple subject sequences were hits for a single query sequence, the ortholog with the highest Blastp score was chosen. Query and subject accession information is provided in Supplemental File 3. **(B)** *S. rosetta* produces chondroitin that can be degraded by ABC chondroitinase and EroS. Polysaccharides isolated from *S. rosetta* were treated with either ABC chondroitinase from *P. vulgaris* (center plot) or EroS (bottom plot). Degradation products from samples treated with ABC chondroitinase and EroS were separated by SAX-HPLC (X-axis indicates time, Y-axis indicates abundance) and compared to the following chondroitin disaccharide standards (top plot): (1) D0a0, unsulfated chondroitin; (2) D0a6, chondroitin-6-sulfate; (3) D0a4, chondroitin-4-sulfate; (4) D0a10, chondroitin-4,6-sulfate; (5) D2a4, chondroitin-2,4-sulfate. Unsulfated and 6-sulfated chondroitin disaccharides were present at similar abundance in both the ABC chondroitinase and EroS –treated samples, whereas all other chondroitin disaccharides were below the limit of detection. **(C)** Quantification of chondroitin disaccharide products produced by *ABC* chondroitinase (*P. vulgaris*) and EroS treatment of *S. rosetta* polysaccharides. Disaccharide abbreviations: D0a0=unsulfated chondroitin; D0a6=chondroitin-6-sulfate; D0a4=chondroitin-4-sulfate; D0a10=chondroitin-4,6-sulfate; D2a4=chondroitin-2,4-sulfate. **(D)** *S. rosetta* does not produce heparan sulfate. Polysaccharides isolated from *S. rosetta* (bottom plot) were treated with Heparinase I, Heparinse II, and Heparinase III (Dextra Laboratories) separated by SAX-HPLC (X-axis indicates time, Y-axis indicates abundance) and compared to the following heparan sulfate disaccharide standards (top plot): (1) D0A0; (2) D0S0; (3) D0A6; (4) D2A0; (5) D0S6; (6) D2S0; (7) D2A6; (8) D2S6. No heparan disaccharides were present above the limit of detection in the *S. rosetta* polysaccharide sample.

**Figure S4.**
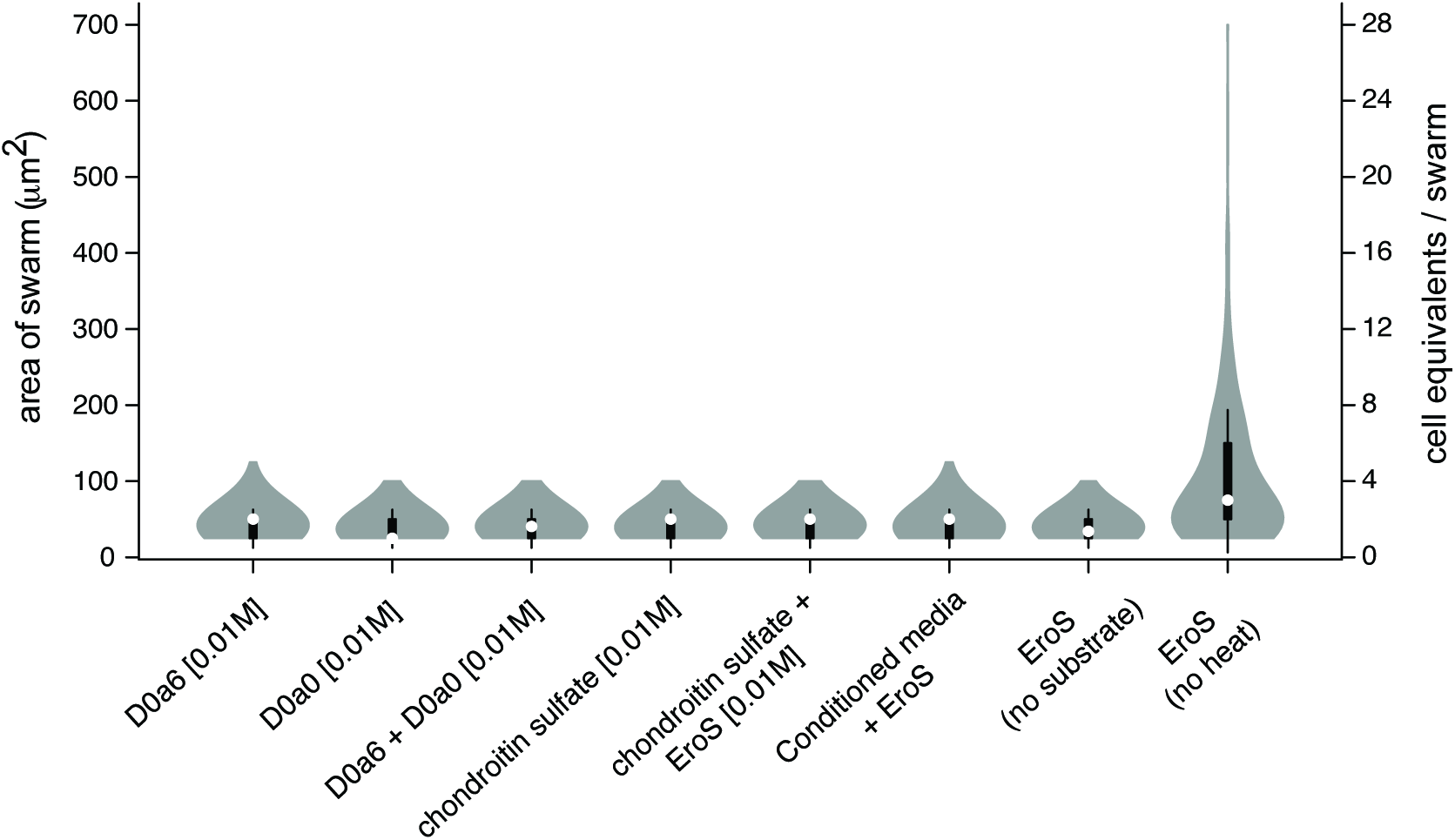
Swarming in *S. rosetta* is not induced by chondroitin sulfate or chondroitin disaccharides. Neither commercial chondroitin disaccharides (DOa6 and DOaO), chondroitin disaccharides generated via the depolymerization of chondroitin sulfate by EroS, nor conditioned media from EroS-treated *S. rosetta* cells are sufficient to induce swarming in *S. rosetta*.

**Figure S5.**
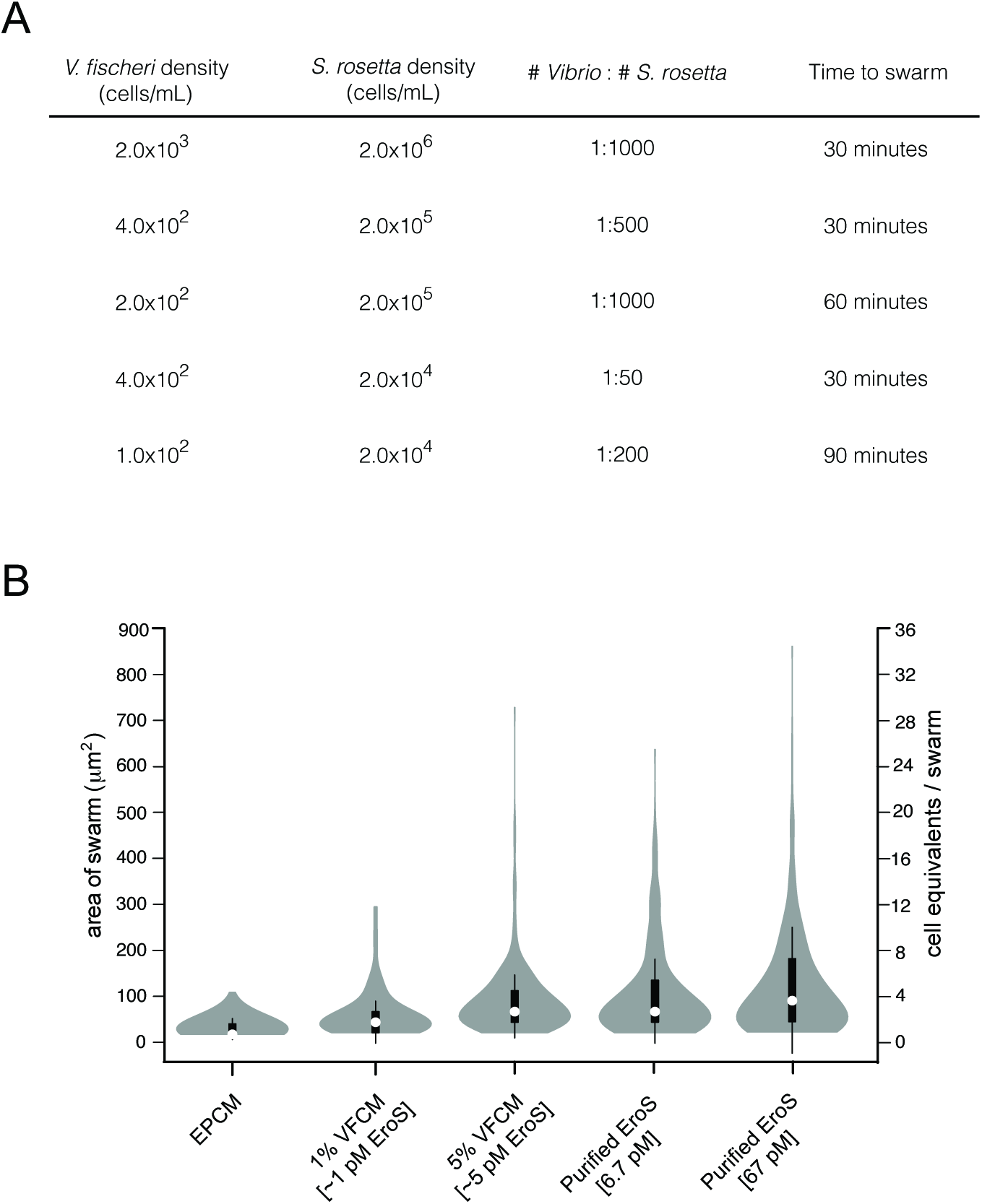
*V. fischeri* induces mating in *S. rosetta* under plausible environmental conditions (related to Table S4). **(A)** *S. rosetta* swarms in response to low numbers of *V. fischeri* bacteria in a cell density-dependent manner. *S. rosetta* at high cell densities (2.0×10^6^ cells/mL) swarms in response to as few as one *V. fischeri* cell per 1000 *S. rosetta* cells within 30 minutes of exposure, whereas swarming in *S. rosetta* at lower cell densities (2.0×10^5^ cells/mL) within a similar time frame requires at least one *V. fischeri* cell per 500 *S. rosetta* cells. **(B)** Picomolar concentrations of secreted (5% VFCM) and purified EroS are sufficient to induce swarming in *S. rosetta*.

**Movie S1 (related to** Figure 1**). Time-lapse movie of *S. rosetta* cells swarming in response to *V. fischeri.*** The movie beings with a side-by-side comparison of *S. rosetta* treated with EPCM (left) and VFCM (right) one hour post induction.

After the short transition (“Induction with *V. fischeri*”) is a time-lapse depicting early swarm formation after treatment with 5% VFCM. All footage is displayed at 35x real time.

**Movie S2 (related to** Figure 1). Two cells within a four-cell swarm undergo cell fusion. Movie begins 30 minutes after addition of 5% VFCM. Cell fusion is displayed at 60X real time.

**Supplemental File 1 (related to** Figure 1). KASP genotyping of meiotic progeny isolated from *V. fischeri*- induced cross

**Supplemental File 2 (related to** Figure 2). Mass Spectrometry of bioactive *V. fischeri* protein fraction

**Supplemental File 3 (related to** Figure 3, Figure S3). Putative homologs of GAG biosynthetic enzymes

## Supplemental Notes

Only 60 validated polymorphisms differentiate *S. rosetta* strains R+ and R-^21^. Therefore, while we are able to detect independent assortment and recombination, because few markers sit within close proximity it was not possible to utilize genotyping data to accurately measure recombination frequency and construct linkage groups.

**Supplemental Table 1.** Bacteria tested in swarming bioassay (related to Figure 2)

| Species | Strain | Genotype | Live bacteria | Conditioned media | Concentrated conditioned media | Accession information |
| --- | --- | --- | --- | --- | --- | --- |
| *Vibrio fischeri* | ES114 | WT |  |  | + | ATCC 700601 |
| | | ΔLuxO | + | + | + | Lupp et al. 2003 |
| | | ΔAinS | + | + | + | Lupp et al. 2003 |
| | | ΔLuxI | + | + | + | Lupp et al. 2003 |
| | | ΔAinSΔLuxS | + | + | + | Lupp and Ruby 2005 |
| | | ΔLuxR | + | + | + | Lupp and Ruby 2005 |
| | | ΔSypC | + | + | + | Lupp and Ruby 2005 |
| | | ΔSypH | + | + | + | Shibata et al. 2012 |
| | | ΔSypI | + | + | + | Shibata et al. 2012 |
| | | ΔSypK | + | + | + | Shibata et al. 2012 |
| | | ΔSypM | + | + | + | Shibata et al. 2012 |
| | | ΔSypO | + | + | + | Shibata et al. 2012 |
| | | ΔSypQ | + | + | + | Shibata et al. 2012 |
|  |  |  | + | + |  | Shibata et al. 2012 |
| *Vibrio fischeri* | MJ11 |  | + | + | + | BAA-1741 |
| *Vibrio tubiashii* |  |  | + | + | + | ATCC 19105 |
| *Vibrio orientalis* |  |  | + | + | + | ATCC 33934 |
| *Vibrio harveyi* |  |  | – | – | – | ATCC 14126 |
| *Vibrio natriegens* |  |  | – | – | – | ATCC 8110 |
| *Vibrio parahaemolyticus* |  |  | ψ | ψ | ψ | ATCC 17802 |
| *Vibrio alginolyticus* |  |  | ψ | ψ | ψ | PRJNA13571 |
| *Vibrio anguillarum* |  |  | – | – | – | ATCC 19181 |
| *Vibrio ordalii* |  |  | – | – | – | ATCC 33509 |
| *Vibrio cholera* | YB1A01 |  | n.t. | – | – | PRJNA281423 |
| *Vibrio metoecus* | YB4D01 |  | – | – | – | PRJNA281423 |
| *Vibrio mimicus* |  |  | – | – | – | ATCC 33653 |
| *Echinicola pacifica* |  |  | – | – | – | DSM 19836 |
| *Echinicola pacifica* | From starved *S. rosetta* co-culture |  | n.t. | – | – | ATCC PRA-390 |

+: swarming observed; –: no swarming observed; ψ: settling observed; n.t.: not tested

**Supplemental Table 2.**
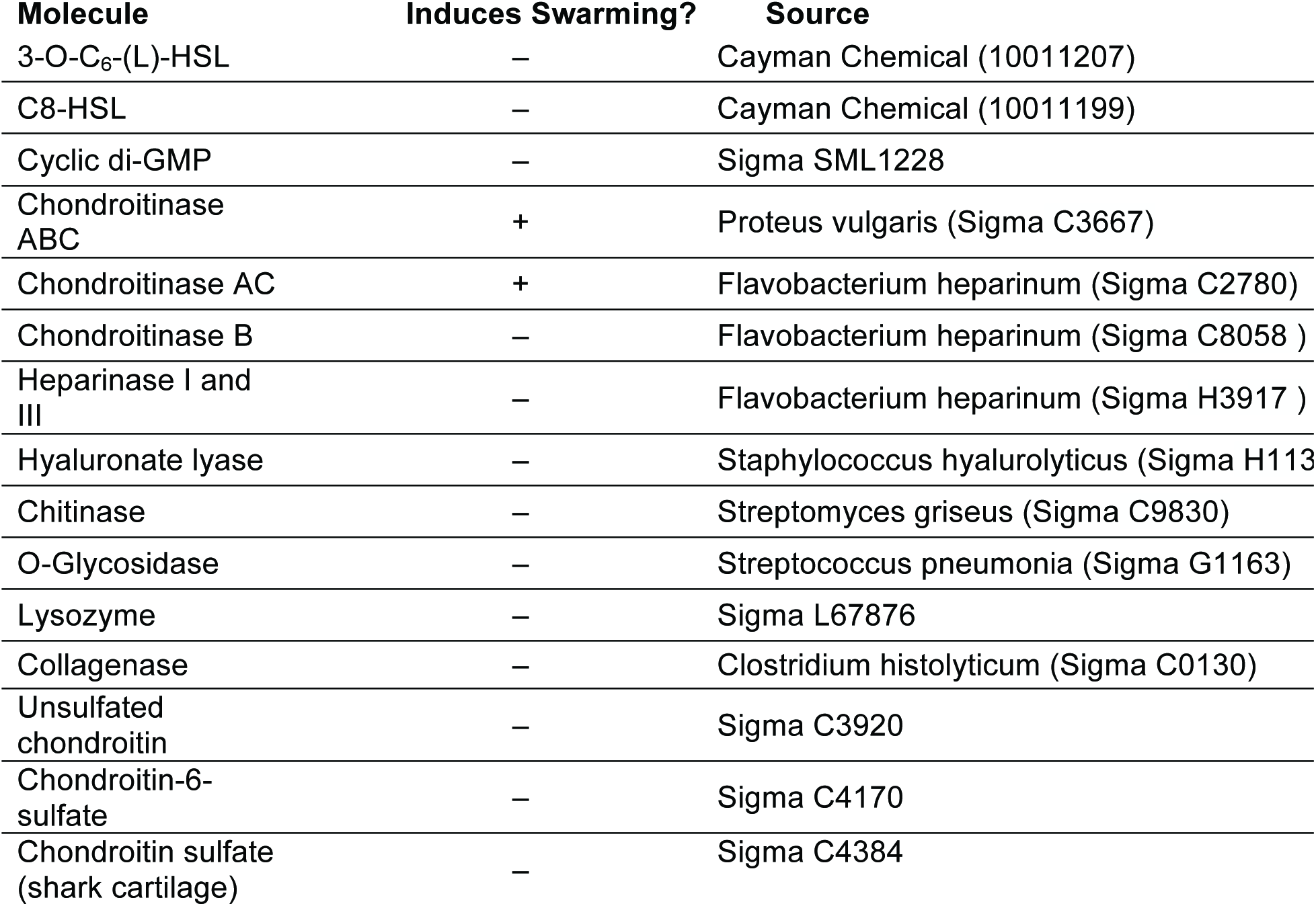
Purified molecules tested in swarming bioassay (related to Figure 2)

| **Molecule** | **Induces Swarming?** | **Source** |
| --- | --- | --- |
| 3-O-C<sub>6</sub>-(L)-HSL | – | Cayman Chemical (10011207) |
| C8-HSL | – | Cayman Chemical (10011199) |
| Cyclic di-GMP | – | Sigma SML1228 |
| Chondroitinase ABC | + | Proteus vulgaris (Sigma C3667) |
| Chondroitinase AC | + | Flavobacterium heparinum (Sigma C2780) |
| Chondroitinase B | – | Flavobacterium heparinum (Sigma C8058 ) |
| Heparinase I and III | – | Flavobacterium heparinum (Sigma H3917 ) |
| Hyaluronate lyase | – | Staphylococcus hyalurolyticus (Sigma H113) |
| Chitinase | – | Streptomyces griseus (Sigma C9830) |
| O-Glycosidase | – | Streptococcus pneumonia (Sigma G1163) |
| Lysozyme | – | Sigma L67876 |
| Collagenase | – | Clostridium histolyticum (Sigma C0130) |
| Un sulfated chondroitin | – | Sigma C3920 |
| Chondroitin-6-sulfate | – | Sigma C4170 |
| Chondroitin sulfate (shark cartilage) | – | Sigma C4384 |

**Supplemental Table 3.** Chondroitinase-induced mating in *S. rosetta* (related to Figure 3)

| Inducing factor | Clonal isolates (#) | % thecate isolates<br>(presumable diploid) | % outcrossed diploid<br>isolates |
| --- | --- | --- | --- |
| *E. pacifica* CM<br>(5% Vol/Vol) | 88 | 0 | 0 |
| EroS<br>(*V. fischeri*, 50pM) | 53 | 17% | 15% |
| Chondroitinase ABC<br>(*P. vulgaris*, 1 unit) | 62 | 21% | 17% |
| Chondroitinase AC<br>(*F. heparinum*, 1 unit) | 52 | 13% | 11% |

**Supplemental Table 4.** Quantification of purified EroS and EroS secreted by *V. fischeri* (related to Supplemental Figure 4)

|  | Purified EroS | Low nutrient media | High nutrient media |
| --- | --- | --- | --- |
| MS2 Peak intensity (sequence TQITDDTYQNFFDK) | 1.43E+05 | 9.08E+02 | 2.77E+03 |
| MS2 Peak intensity (1 pMol standard) | 1.07E+04 | 2.24E+04 | 1.86E+04 |
| pMol EroS in excised gel band | 13.4 | 0.0405 | 0.149 |
| Sample volume in gel band (μL) | 2.0 | 18.0 | 6.0 |
| [EroS] loaded on gel | 6.7 μM | 2.3 nM | 25 nM |
| [EroS] in culture supernatant |  | 6.8 pM | 75 pM |

